# Comparative transcriptional analysis reveals gene expression changes in spore germination of opportunistic pathogenic fungi

**DOI:** 10.1101/2025.02.03.636182

**Authors:** Da-Woon Kim, Malaika K. Ebert, Soumya Moonjely, Zheng Wang, Oded Yarden, Jeffrey P. Townsend, Frances Trail

**Affiliations:** Department of Plant Biology, Michigan State University, East Lansing, Michigan, USA 48854; Department of Plant Pathology, North Dakota State University, Fargo, North Dakota, USA; Department of Biostatistics, Yale School of Public Health, New Haven, Connecticut, USA; Department of Plant Pathology and Microbiology, The Robert H. Smith Faculty of Agriculture, Food and Environment, The Hebrew University of Jerusalem, Rehovot, Israel; Department of Plant, Soil and Microbial Sciences, Michigan State University, East Lansing, Michigan, USA 48854

## Abstract

In opportunistic human pathogenic fungi, changes in gene expression play a crucial role in the progression of growth stages from early spore germination through host infection. Comparative transcriptomics between diverse fungal pathogens and non-pathogens provided insights into regulatory mechanisms behind the initiation of infectious processes. We examined the gene expression patterns of 3,845 single-copy orthologous genes (SCOGs) across five phylogenetically distinct species, including the opportunistic human pathogens *Fusarium oxysporum*, *Aspergillus fumigatus*, and *A. nidulans,* and nonpathogenic species *Neurospora crassa* and *Trichoderma asperelloides*, at four sequential stages of spore germination. Ancestral status of gene expression was inferred for nodes along the phylogeny. By comparing expression patterns of the SCOGs with their most recent common ancestor (MRCA), we identified genes that exhibit divergent levels of expression during spore germination when comparing fungal pathogens to non-pathogens. We focused on genes related to the MAPK pathway, nitrogen metabolism, asexual development, G-protein signaling, and conidial-wall integrity. Notably, orthologs of the transcription activator *abaA*, a known central regulator of conidiation, exhibited significant divergence in gene expression in *F. oxysporum*. This dramatic expression change in *abaA* was accompanied by structural modifications of phialides in *F. oxysporum*, and revealed how these changes impact development of offspring, formation of aerial hyphae, spore production, and pathogenicity. Our research provides insights into ecological adaptations observed during the divergence of these species, specifically highlighting how divergence in gene expression during spore germination contributes to their ability to thrive in distinct environments.

**Author Summary:** The fungi of the phylum Ascomycota include plant and animal pathogens, endophytes, and saprotrophs, some of which are model organisms for biological investigation, and usually have abundant genomic and transcriptomic data. In this study, transcriptomics was studied during spore germination in five species: the opportunistic human pathogens *Fusarium oxysporum*, *Aspergillus fumigatus*, and *Aspergillus nidulans*, and the nonpathogenic species *Neurospora crassa* and *Trichoderma asperelloides*. We have inferred divergence in gene expression for 3,845 single-copy orthologous genes (SCOGs) along these lineages. Genes related to the MAPK pathway, nitrogen metabolism, conidia-related regulators, G protein signaling, and conidial-wall integrity exhibited dramatic expression shifts in the lineages of the opportunistic human pathogens evolving towards true pathogenic species. Notably, *abaA*, a known central regulator of conidiation, exhibited striking divergence in expression in the opportunistic pathogens, resulting in structural modifications in phialides that were more similar to pathogenic species, and revealing how these changes impact development of aerial hyphal formation, spore production, and pathogenicity. These findings provide insights into ecological adaptations resulting from divergence in gene expression, and reveal dynamic transcriptional changes, which may be crucial for the adaptation of opportunistic pathogens to changing environments. By elucidating the shifting roles of *abaA*, our research contributes to a deeper understanding of the divergence mechanisms underlying development and pathogenicity in fungi.

## Introduction

Spore germination plays a critical role in the life cycle of fungi, facilitating their propagation and spread by transitioning them from a dormant state to active growth (1–4). This transition is marked by a series of adaptive responses, as germinating spores face and overcome environmental challenges such as nutrient scarcity, osmotic stress, and exposure to biotic and abiotic factors (5–10). Identifying the molecular mechanisms underlying spore germination and colony initiation is essential for understanding the establishment and spread of fungal diseases, as well as for developing control strategies (11–13).

Previous studies exploring transcriptional changes during spore germination in fungal species have uncovered key regulators and signaling pathways involved in the process (14–16). Our previous work demonstrated that integrating comparative genomics, transcriptomics, and gene knockout techniques within a phylogenetic framework efficiently identified genes whose shifts in expression drive developmental and phenotypic divergence among species (17–19). This comparative approach revealed key genes with roles in sexual development of *N. crassa* and *F. graminearum* (17). Transcriptional divergence through evolutionary processes was shown to be crucial for multicellular fruiting body development, with genes encoding hypothetical proteins playing essential roles in *F. graminearum*, *M. oryzae*, and *N. crassa* (18). Comparative transcriptomics of spore germination in *F. graminearum* and *M. oryzae* revealed novel insights into the plant infection process (19). These findings emphasize the critical role transcriptional divergence plays in establishment of morphological and functional traits, including in sexual development and spore germination.

Here, we illuminate the differences in gene expression patterns during spore germination between opportunistic human pathogenic fungi and non-pathogenic fungi. We examine transcriptional divergence in cellular processes related to spore germination, growth, survival, and pathogenicity, by focusing on single-copy orthologous genes (SCOGs) among five species. To identify expression changes unique to opportunistic human pathogens, we analyzed and inferred the ancestral transcriptional profiles of these species, during spore germination, from divergent lineages in the Ascomycota (**Fig. 1A**). *A. fumigatus* is a major pathogen responsible for aspergillosis in individuals with compromised immune systems (20). The *F. oxysporum* isolate used here is an opportunistic human pathogen isolated from a leukemia patient (21). *A. nidulans* is a model organism for cellular and molecular biology research, and occasionally leads to ocular infections (22). *N. crassa* serves as a key model organism for research in genetics and biochemistry (23, 24). *T. asperelloides* is known for its biocontrol and plant growth promotion properties (25). We demonstrate the unique and emergent gene expression patterns during spore germination in these fungi ranging from opportunistic human pathogens to saprotrophs, through a comparative transcriptomic analysis. This comprehensive investigation infers genes and pathways critical for spore germination and elucidates the pathogenic function of the *abaA* gene, which exhibits a unique expression pattern in the opportunistic human pathogen *F. oxysporum*.

**Figure 1.**
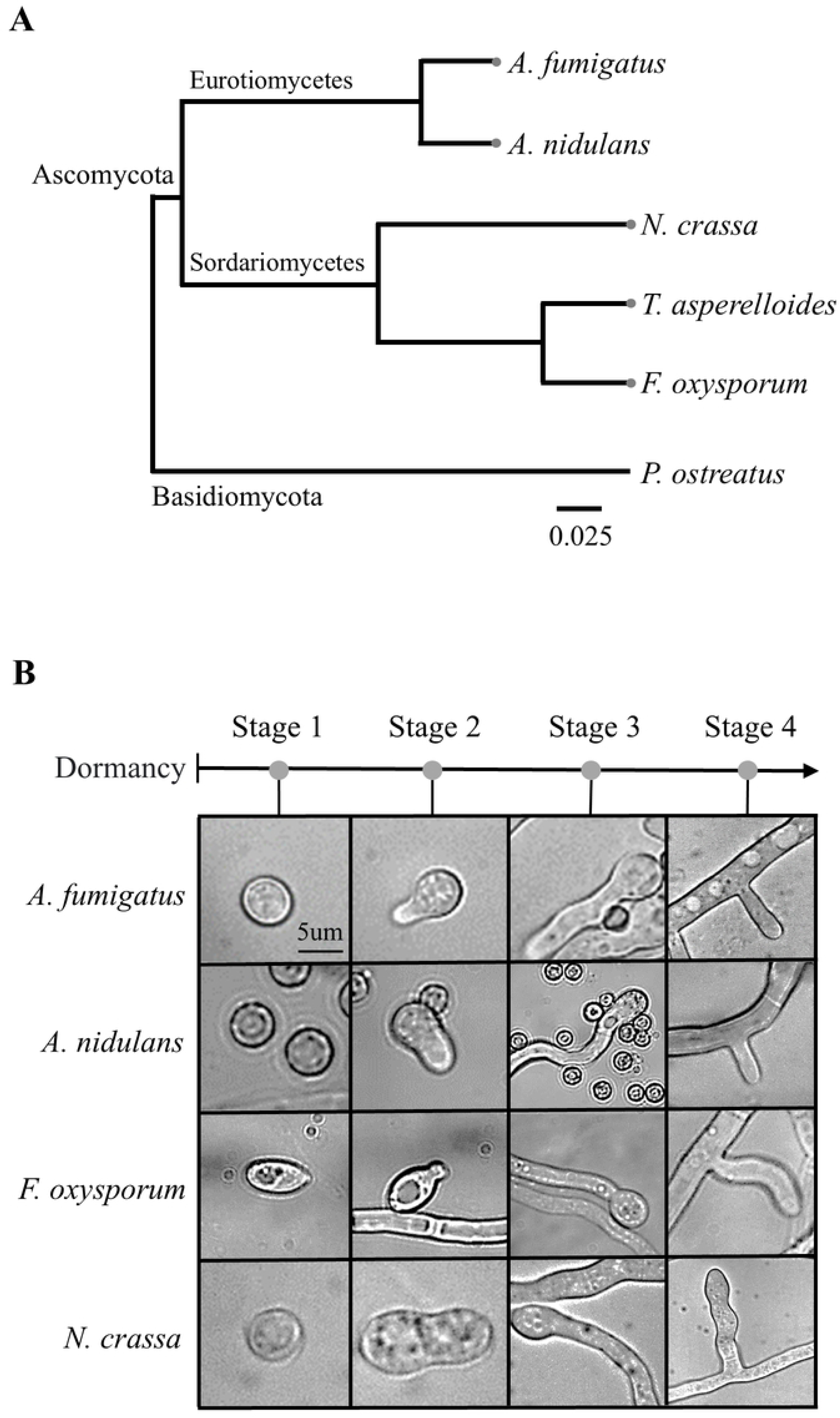

## Results

### Transcriptome analysis of divergence in expression during four stages of spore germination

Conidia of *F. oxysporum*, *A. nidulans*, and *A. fumigatus* presented the four defined stages of spore germination (**Fig. 1B**) and the transitions from stage 1 to stage 4 were completed within 12 hours. The RNA-sequencing data, processed using Htseq-count, showed 85.56-99.49% mapping rates across all stages (**Table S1**). Transcriptional divergence was analyzed for the 3,845 single copied orthologous genes (SCOGs) conserved in the five species, with *N. crassa* and *T. asperelloides* serving as two nonpathogenic references (**Table S2**). During axis extension (Stages 2 to 3), the fold change differential (FCD) values exhibited a relatively small range, from −41.13 to 18.27. In contrast, large changes in the expression patterns of genes were identified during the process of germ tube formation (Stages 1 to 2) and hyphal branching (Stages 3 to 4) (**Table S3**). The FCD values of the conserved genes *EF-1* and *RPB2* (referred to as housekeeping genes) ranged from −2 to 2 across all stages for all species (**Fig. S1B**). This consistent range confirmed that there were no significant changes in the expression of the housekeeping genes. This analysis highlights key transcriptional divergences during spore germination, setting the stage for a more detailed exploration of the functional groups involved in these evolutionary changes.

### *HamE,* the mitogen-activated protein kinase (MAPK) complex scaffolding protein, exhibits highly divergent expression among species

*HamE*, a MAPK complex scaffolding protein, exhibits highly divergent expression among the species. In *A. nidulans, HamE* (AN2701) positively regulates the phosphorylation of the MAP kinase *MpkB* (AN2393), resulting in increased expression of genes involved in development and secondary metabolism (38–41). We analyzed the MAPK pathway network based on the KEGG pathway and constructed a schematic diagram incorporating *HamE* (**Fig. 2A**). This MAPK pathway, consisting of 12 genes, is well-conserved across the five species examined, except for orthologs of *Ste2*, *Ste3*, and *LaeA*. The SCOGs *HamE*, *FlbA*, and *MpkB* exhibited expression variation in fold change (FC) values across stages among the species (**Fig. 2B**). To identify the most significantly expressed genes, we applied criteria based on the top 10% ranking and FC > 2, which indicated that *MpkB* (NCU02393) was highly expressed during germ tube formation in *N. crassa*, showed a FC of 14.7 and ranked 151st compared to the 3,845 SCOGs in the five species. Additionally, *HamE* (AN2701) showed divergence in expression during hyphal branching in *A. nidulans* (FC 43.8, ranked 3rd), and during axis extension in both *A. fumigatus* (FC 14.1, ranked 1st) and *T. asperelloides* (FC 4.9, ranked 73rd). Conversely, when applying criteria based on the bottom 10% ranking and FC < −2, *FlbA* (NCU08319) showed expression changes indicating decreased expression levels during germination in *N. crassa* (FC −5.4, ranked 3759th), and *HamE* (AFUA_5G13970) also exhibited decreased expression during hyphal branching in *A. fumigatus* (FC −3.1, ranked 3837th). These results demonstrate the divergence of gene expression within the pheromone module, and the comparatively high expression of *HamE* in *A. fumigatus* and *A. nidulans* relative to other species (**Fig. 2B**). These findings suggest that *HamE* plays a critical role in development and stress response processes.

**Figure 2.**
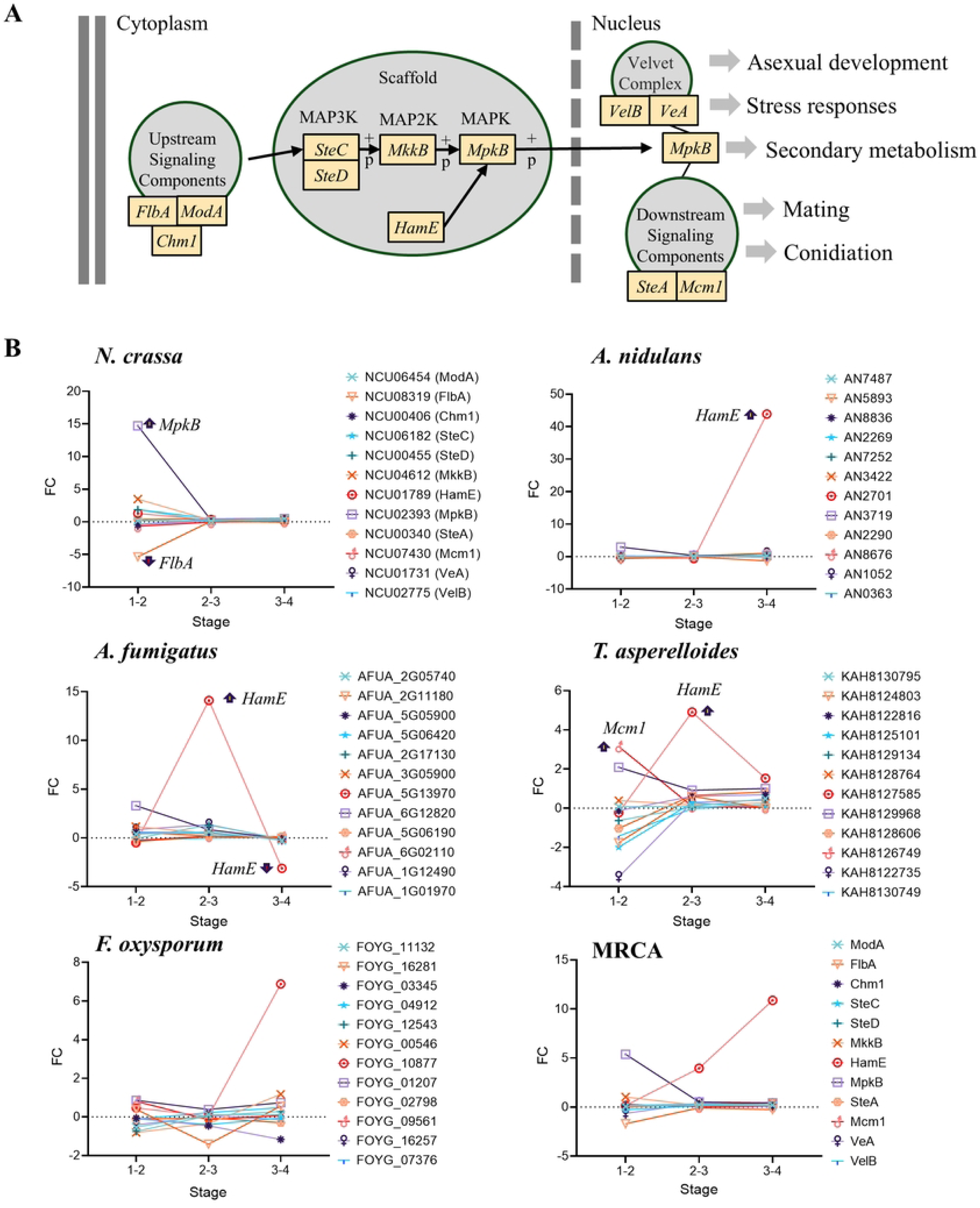

### Comparative analysis of gene expression for nitrogen metabolism during spore germination

We examined the expression of 16 nitrogen metabolism-related genes in *A. nidulans* identified via the KEGG pathway, with their single-copy orthologs (SCOGs) found to be well-conserved among the other four species (**Fig. S2A, B**). During spore germination, minimal changes were observed in the expression of these genes in *A. nidulans* and *A. fumigatus*. Specifically, only one nitrogen metabolism-related gene ranked within the top 10% (AN1006) and bottom 10% (AFUA_1G10930) in expression levels in each of these two species, suggesting limited transcriptional activity in this group during germination. In contrast, *F. oxysporum* harbors eight nitrogen metabolism-related genes ranked within the top or bottom 10%, whereas *N. crassa* and *T. asperelloides* each had five genes that were either up- or downregulated (**Table S4**). This comparison highlights the disparity between the *Aspergillus* species, which had only one gene ranked, and *F. oxysporum*, *N. crassa*, and *T. asperelloides*, which exhibited more pronounced transcriptional responses.

### Key gene expression changes in asexual development

Previous studies have shown that conidiation is modulated by upstream regulators (*FluG*, *FlbA*, *FlbB*, *FlbC*, *FlbD*, and *FlbE*), negative regulators (*SfgA*, *StuA*, and *MedA*), velvet regulators (*VeA*, *VelC*, *VelB*, and *VosA*), and central regulators (*BrlA*, *AbaA*, and *WetA*) (26–28) (**Fig. 3A**). Divergent regulation of gene expression was observed across the species examined. In *A. nidulans*, *EsdC* (AN9121) (29) and *StuA* (AN5836) exhibited highly divergent changes in gene expression during developmental processes. Similarly, in *A. fumigatus*, both *EsdC* (AFUA_7G01930) and *StuA* (AFUA_2G07900) showed highly divergent changes in their expression patterns (**Tables S4, and S5**). While the two *Aspergillus* species demonstrated similarities in the expression profiles of *EsdC* and *StuA*, such as the presence of highly divergent changes in both species, the timing of their expression ranking differed. Additionally, *StuA* exhibited expression changes that were specific to *Aspergillus* during spore germination, while no ranked expression changes in *StuA* orthologs were observed in non-Aspergillus species at the same developmental stages (**Fig. 3B**). In *F. oxysporum*, *EsdC* (FOYG_08090) ranked 3734th during germination and ranked 24th during hyphal branching. *AbaA* (FOYG_06458) ranked 3825th during hyphal branching, showing a *F. oxysporum*-specific pattern (**Fig. 3B**). In *T. asperelloides*, *EsdC* (KAH8130575) ranked 3713th during germination and ranked 4th during axis extension. *FlbC* (KAH8128436) ranked 3824th during germination, showing a *T. asperelloides* specific pattern. In *N. crassa*, *EsdC* (NCU03600) ranked 48th, *PkaA* (NCU06240) ranked 66th, and *FlbA* (NCU08319) ranked 3759th during germination, and *PkaA* (NCU06240) and *FlbA* (NCU08319) exhibited *N. crassa* specific expression patterns (**Fig. 3B**; **Table S5**).

**Figure 3.**
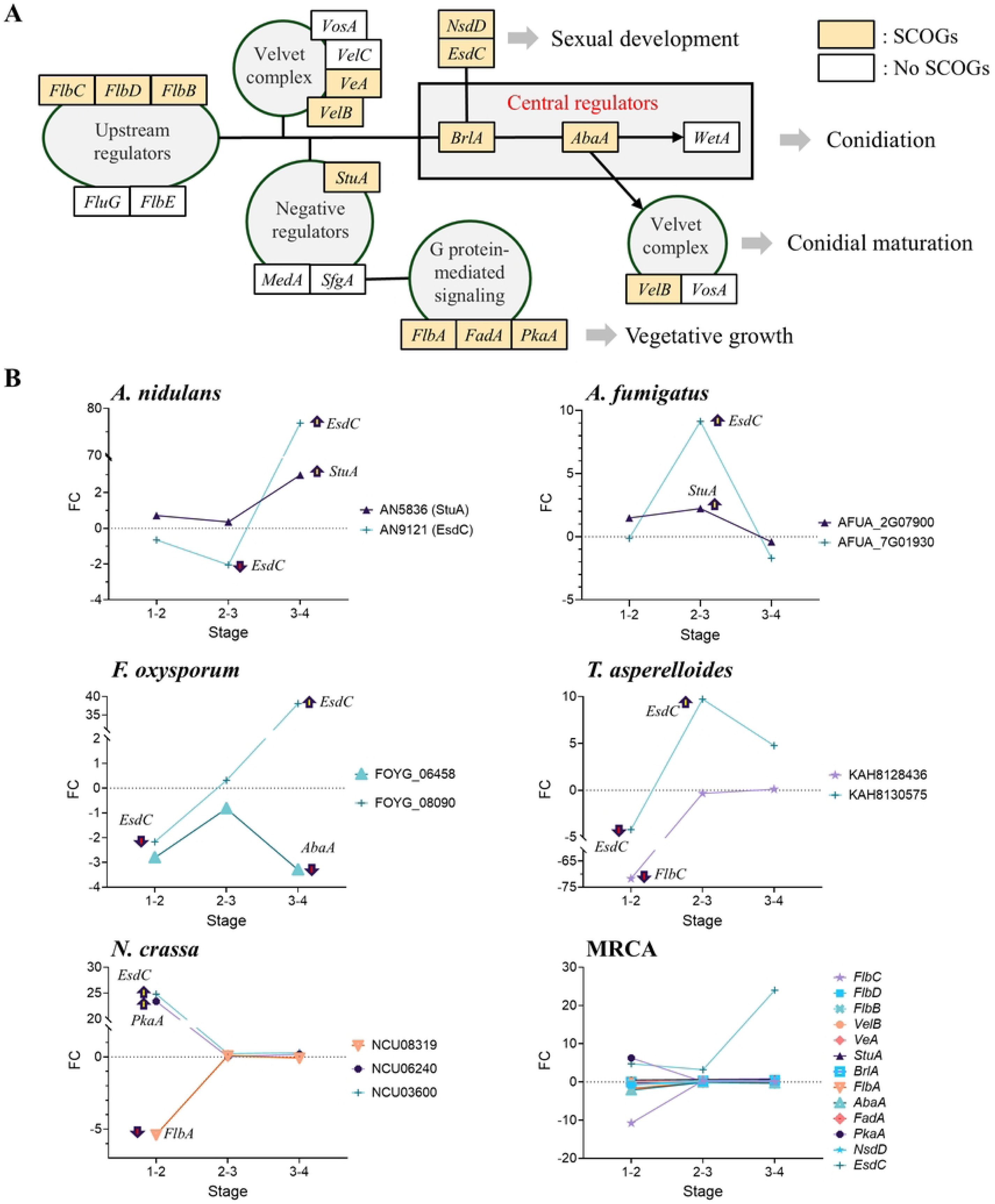

### Transcriptional divergence during spore germination

In the opportunistic human pathogens *A. nidulans*, *A. fumigatus*, and *F. oxysporum*, several genes exhibited specific expression changes (**Tables S4, and S5**). *GprK,* a regulator of G protein signaling with an RGS domain, showed a distinct expression pattern compared to non-pathogenic species, ranking highly during spore germination in the opportunistic human pathogens *A. fumigatus* (AFUA_4G01350), *A. nidulans* (AN7795), and *F. oxysporum* (FOYG_11952) (**Tables S4, and S5**). During spore germination, *BglA*, involved in conidial-wall integrity, was top ranked in *A. fumigatus* (AFUA_1G05770; 81st) and *A. nidulans* (AN4102; 68th) compared to *F. oxysporum* (FOYG_02502; 2933rd), *T. asperelloides* (KAH8126432; 2910th), and *N. crassa* (NCU08755; 3828th) (**Tables S4, and S5**). These findings suggest that *BglA* and *GprK* exhibit notable divergence in gene expression in pathogens compared to non-pathogens, indicating that changes in the cell wall, or in the regulation of G protein signaling, play a role in stress resistance and host adaptation. *SidD* and *HapX* are involved in iron homeostasis, and in this category, *SidD* (AFUA_3G03420) ranked 111th during germination in *A. fumigatus*, and *HapX* (AN8251) ranked 3465th during germination in *A. nidulans*. *Sod2* (FOYG_12102), a gene involved in stress response, is highly ranked, at 138th, during hyphal branching exclusively in *F. oxysporum*, indicating its significant role in that species compared to the other. In addition to these genes, our analysis revealed significant shifts in gene expression across various stages of fungal development, including notable changes in the expression of four hypothetical proteins (**Table S5**). We have highlighted specific genes, including *GprK*, *SidD*, *HapX*, and *Sod2*, that exhibited distinct, stage-specific expression changes during spore germination in the opportunistic human pathogens *A. nidulans*, *A. fumigatus*, and *F. oxysporum*, suggesting their potential roles in pathogen-specific adaptations during spore germination.

### Divergence of *abaA* expression related to conidiation and pathogenicity

To investigate the divergence in the expression of *abaA* and its role in conidiation and pathogenicity, we studied *abaA* from *F. oxysporum* (FOYG_06458; *FoabaA*) and from *A. fumigatus* (AFUA_1G04830; *AfabaA*). We compared the promoter regions of *FoabaA* and *AfabaA*, finding that P_FoabaA_ included the 872 bp 5’ UTR, while P_AfabaA_ had the 979 bp 5’ UTR, with 40.49% sequence similarity (**Fig. S4, Table S6**). BLAST analysis of the predicted protein sequences of *FoabaA* and *AfabaA* showed 10% query cover, 42% identity, and 19% gaps (**Table S7**). Integration into the mutants was confirmed through Sanger sequencing and revealed nucleotide sequence identities of 99.52% to 99.86% for the promoters and genes, and protein sequence identities of 99.12% to 99.87% (**Table S8**). Growth observations on CM, PDA, and Bird agar medium revealed that Δ*FoabaA* exhibited increased aerial hyphae on CM and PDA media compared to WT, with no differences on Bird medium (**Fig. 4**). Additionally, the Δ*FoabaA*:: P_AfabaA_:: *AfabaA* strain did not produce pigment, unlike WT, and the Δ*FoabaA*:: P_FoabaA_:: *AfabaA* strain showed reduced aerial hyphae. Mutants containing *AfabaA* showed weaker aerial hyphae formation compared to those with *FoabaA* (**Fig. 4**). Expression levels of *abaA* in each strain were categorized based on the promoter of *FoabaA* or *AfabaA* as follows: WT (native expression, NE::*FoabaA*), Δ*FoabaA*:: P_AfabaA_:: *AfabaA* (overexpression, OE::*AfabaA*), Δ*FoabaA*:: P_AfabaA_:: *FoabaA* (OE::*FoabaA*), and Δ*FoabaA*:: P_FoabaA_:: *AfabaA* (NE::*AfabaA*). The expression of OE::*FoabaA* and OE::*AfabaA* strains were validated through qRT-PCR (**Fig. 5B**). qRT-PCR results indicated an increase in the FC for *AfabaA* with its promoter in the axis extension and hyphal branching compared to the native promoter of *FoabaA*. Phenotypic characteristics, including germination, conidiation, pathogenicity, and phialide morphology, are summarized in **Table 1**. Both Δ*FoabaA* and NE::*AfabaA* strains failed to produce spores and formed elongated phialides (**Fig. S3**). The OE::*AfabaA* and OE::*FoabaA* strains showed reduced conidiation, displaying a mixture of WT-like and elongated phialides, with the OE::*FoabaA* strain showing chain-type phialides (**Figs. 5A, S3**). Pathogenicity tests on *G. mellonella* showed faster disease progression in both OE strains compared to WT, advancing by approximately one day (**Fig. 5C**).

**Figure 4.**
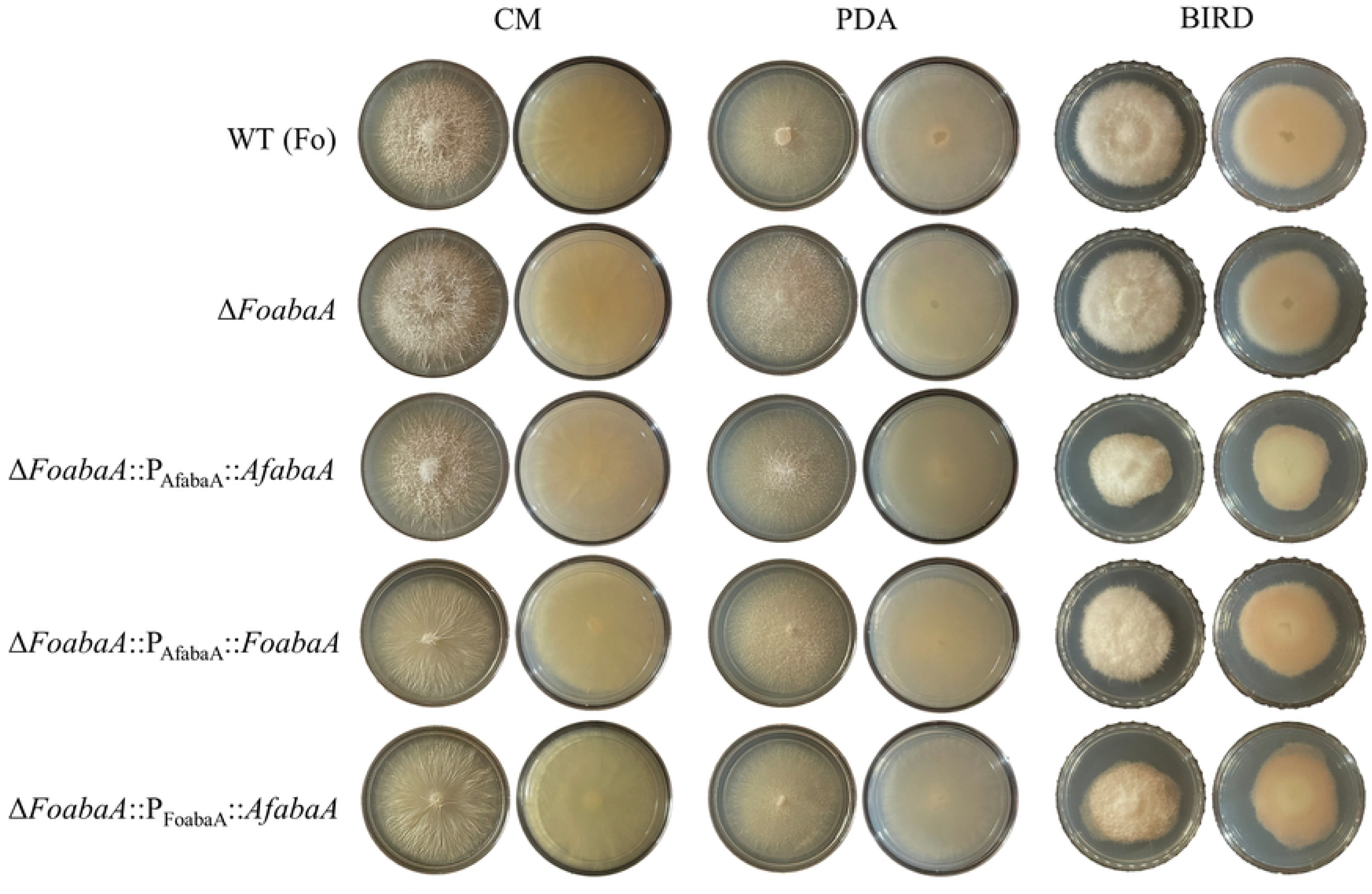

**Figure 5.**
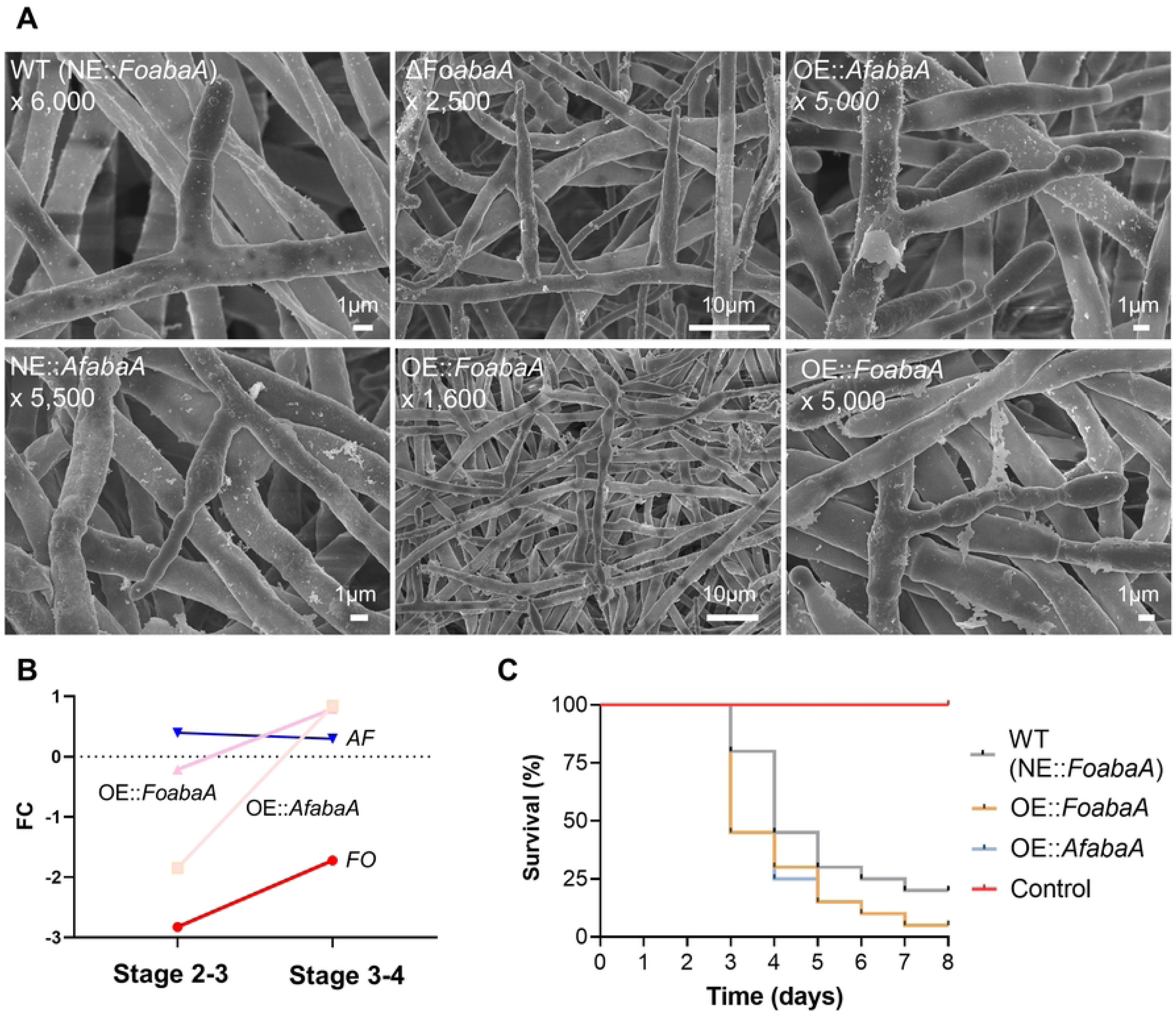

**Table 1.** Summary of phenotypes of *abaA*-related mutant strains.

| STRAIN | Germination | Conidiation | Pathogenicity | Morphology of Phialides |
| --- | --- | --- | --- | --- |
| NE::*FoabaA*<br>(WT) | WT | WT | WT | WT |
| Δ*FoabaA* | No conidia | No conidia | No conidia | Elongated phialides |
| OE::*AfabaA* | WT | Decreased | Increased | WT;<br>Elongated phialides |
| OE::*FoabaA* | WT | Decreased | Increased | WT;<br>Elongated phialides;<br>Chain form |
| NE::*AfabaA* | No conidia | No conidia | No conidia | Elongated phialides |

### Transcriptome analysis of overexpressed *FoabaA* and *AfabaA* in *F. oxysporum*

We performed transcriptome analysis to investigate the divergence in gene expression of WT vs OE::*AfabaA* and WT vs OE::*FoabaA.* DEG analysis was used to investigate the differences in gene expression among WT, OE::*AfabaA*, and OE::*FoabaA*, while Principal Component Analysis (PCA) was used to assess the overall transcriptomic differences among these samples (**Fig. S5**). DEGs were identified using FC values of 1 and higher (up-regulated) or −1 and lower (down-regulated) in comparison to WT (**Fig. S6 and Table S9**). In the OE::*AfabaA* strain, 225 genes were up-regulated and 452 genes were down-regulated. In the OE::*FoabaA* strain, 110 genes were up-regulated and 313 genes were down-regulated. Among these, 74 genes were up-regulated in both OE::*AfabaA* and OE::*FoabaA* strains, representing 32.9% and 67.3% of the up-regulated genes, respectively. Additionally, the same set of 202 genes was down-regulated in each mutant, corresponding to 44.7% and 64.5% of the down-regulated genes, respectively.

### Genes upregulated by overexpression of *abaA* located on the Human Pathogen Lineage Specific chromosomes

We confirmed the chromosomal locations of the upregulated DEGs and found that among the 225 DEGs in the OE::*AfabaA* strain, 110 genes (48.9%) were located on the Human Pathogen Lineage Specific (HPLS) chromosome (**Fig. S6**). Similarly, for the OE::*FoabaA* strain, 40 of 110 upregulated DEGs (36.4%) mapped to the HPLS chromosome. Additionally, 40 of the 74 DEGs (54.1%) upregulated in both OE::*AfabaA* and OE::*FoabaA* strains were located on the HPLS chromosomes (**Fig. S6**). Chromosomal mapping for the up-regulated DEG set is shown in **Fig. S7**. We performed Pfam domain analysis and orthology searches for the 43 genes mapped to the HPLS chromosomes that were upregulated in both OE::*AfabaA* and OE::*FoabaA* strains (**Fig. S8**). Pfam domain analysis revealed the presence of domains commonly found in protein kinases, the glycosyl hydrolases family, the major facilitator superfamily, and cytochrome P450. However, 27 of these genes were not annotated in the Pfam domain analysis. To infer the origins of these 43 genes, we performed an orthology search across seven fungal genera (*Aspergillus*, *Candida*, *Coccidioides*, *Fusarium*, *Histoplasma*, *Neurospora*, and *Trichoderma*), five phyla of the Eukaryota (Metazoa, Alveolines, Amoebozoa, Euglenozoa, Viridiplantae), along with additional phyla classified as ‘others’, and seven strains in the species (*A. fumigatus* Af293, *A. nidulans* A4, *N. crassa* OR74A, *F. oxysporum* f. sp. conglutinans Fo5176, *F. oxysporum* f. sp. cubense, *F. oxysporum* f. sp. lycopersici 4287, and *F. oxysporum* f. sp. melonis 26406) (**Fig. S8**). Eight of these genes were orphan genes with no orthologs, while 14 genes had orthologs only among four plant pathogenic *F. oxysporum* strains. Orthologs of the remaining genes showed widespread distribution across genera of fungi (**Fig. S8**).

## Discussion

Spore germination is a pivotal stage in the fungal life cycle, marking the transition of dormant spores into actively growing states, enabling fungi to colonize environments and establish infections in hosts. Our study reveals significant transcriptional differentiation during germination across the five fungal species through comparative transcriptomic analysis, offering insights into the diversity of gene expression during this process in opportunistic human pathogens and nonpathogens. The transcriptomic profiles revealed stage-specific differentiation among the five species. Notable transitions were observed between spore swelling and germ tube formation (Stage 1-2), and between germ tube elongation and hyphal branching (Stage 3-4), marking critical developmental checkpoints that likely offer survival advantages in ecological or host-associated niches. Our findings reveal transcriptional divergence in the opportunistic human pathogens *A. fumigatus*, *A. nidulans*, and *F. oxysporum*, enabling them to overcome host-associated stresses. For example, differential expression of genes related to cell wall remodeling (*BglA*), stress response (*Sod2*), and iron homeostasis (*SidD* and *HapX*) indicates their role in countering host-derived challenges such as oxidative stress and nutrient limitation (30–32). These findings align with previous studies emphasizing the significance of transcriptional flexibility in fungal pathogenicity and stress resistance (33–38). The MAPK pathway analysis reveals interspecies variation in key regulators including *HamE* and *FlbA*. The differential expression of *HamE* between *A. nidulans* and *A. fumigatus* suggests context-dependent roles in regulating development and secondary metabolism. These findings are consistent with earlier reports that scaffold proteins such as *HamE* modulate MAPK signaling to mediate species-specific developmental processes (39, 40). Nitrogen metabolism is crucial for opportunistic human pathogens, providing essential nutrients for survival and proliferation within the host, thus influencing pathogenicity (41–44). Genes related to nitrogen metabolism exhibit distinct expression patterns, with *F. oxysporum* showing the highest transcriptional activity, possibly reflecting adaptation to ecological niches, including host tissues with variable nitrogen availability. The limited variation in expression of nitrogen metabolism genes in *A. fumigatus* and *A. nidulans* during germination may indicate stable metabolic strategies optimized for their environments. Pathogenicity-related genes, such as the G-protein signaling regulator *GprK*, also exhibited unique expression patterns. The higher expression of *GprK* in these pathogens compared to nonpathogenic species supports its role in mediating environmental sensing and stress resistance, critical for host colonization (45, 46).

*abaA* displayed a species-specific expression pattern in *F. oxysporum*, suggesting a unique role in hyphal branching and pathogenicity. The transcriptional divergence of *abaA* during hyphal branching in *F. oxysporum* suggests the involvement of alternative regulatory pathways in conidiation. This hypothesis aligns with existing studies in *N. crassa* (47), where conidiation persists in *abaA* deletion mutants, demonstrating evolutionary divergence in *abaA* function among fungal species. Furthermore, increased germination following the introduction of *A. nidulans abaA* into *H. capsulatum* underscores functional diversification of *abaA* across the phylum Ascomycota (48). This study experimentally validated the hypothesis of evolutionary changes in *abaA* through promoter and gene swap experiments. Mutants such as Δ*FoabaA*, OE::*AfabaA*, and OE::*FoabaA* revealed critical changes in conidiation and phialide formation. Notably, Δ*FoabaA* and NE::*AfabaA* mutants failed to produce spores, whereas OE::*AfabaA* and OE::*FoabaA* partially restored conidiation with reduced spore production to these three mutants compared to the wild-type (NE::*FoabaA*). Morphological changes, such as elongated phialides in Δ*FoabaA* and NE::*AfabaA* mutants and chain-like phialides in OE::*FoabaA*, further indicated that *abaA* expression levels influence both developmental morphology and reproductive capacity. Pathogenicity assays using *Galleria mellonella* demonstrated rapid pathogenicity in OE::*AfabaA* and OE::*FoabaA* mutants compared to WT, emphasizing the crucial role of *abaA* in fungal pathogenicity. Transcriptomic analysis revealed that 58.1% of the upregulated genes in OE::*AfabaA* and OE::*FoabaA* mutants were located on the HPLS chromosome, which is known to mediate host-specific interactions in plant-pathogenic *F. oxysporum* (21). These findings suggest that transcriptional divergence in *abaA* contributes to pathogenic traits and ecological specialization. Notably, replacing the *FoabaA* promoter with the *AfabaA* promoter mimicked *AfabaA* expression patterns, reducing conidiation and forming Aspergillus-like phialide structures while increasing pathogenicity. These results underscore the evolutionary impact of regulatory mechanisms in *abaA* and the role of *abaA* in adaptation to diverse environments.

*F. oxysporum* NRRL 32931 is classified within the Fusarium oxysporum species complex (FOSC), which encompasses both plant and human pathogens (49). Despite being an opportunistic human pathogen, NRRL 32931 shares a significant portion of its core chromosomes with the plant-pathogenic *F. oxysporum* Fol4287, excluding the HPLS chromosomes (21). The presence of the HPLS chromosomes in NRRL 32931 suggests an adaptive mechanism that enhances its ability to infect and thrive in human hosts, potentially through horizontal gene or chromosome transfer (HGT/HCT) from plant-pathogenic strains in shared environments, such as soil and colonized plants (50). These genetic exchanges may drive the evolution of HPLS chromosomes, facilitating adaptation to diverse environments and enabling pathogenicity in both plant and human hosts.

The transcriptional divergence observed in this study underscores the role of evolutionary pressures in fungal development and pathogenicity. Species-specific expression changes in stress response and nutrient metabolism genes point to genetic innovations that enhance host interactions and environmental resilience. Moreover, transcriptional regulation of *abaA* and associated genes on the HPLS chromosomes contributes to ecological and functional diversification, linking these changes to fungal pathogenicity and adaptive mechanisms. Future research will focus on uncovering the functional roles of dynamically regulated genes to provide deeper insights into the mechanisms of pathogenesis in opportunistic human pathogens. Comparative analyses across fungal pathogens are likely to reveal conserved and divergent evolutionary patterns, ultimately advancing fungal disease management strategies.

## Materials and Methods

### Fungal strains, RNA isolation and RNA sequencing

The genome sequences of strains of five fungal species were used for this study: *A. fumigatus* AF293 (NCBI genome assembly accession no. GCA_000002655.1), *A. nidulans* A4 (NCBI genome assembly accession no. GCA_000149205.2), *F. oxysporum* NRRL 32931 (NCBI genome assembly accession no. GCA_000271745.2), *N. crassa* OR74A (NCBI genome assembly accession no. GCA_000182925.2), and *T. asperelloides* T203 (NCBI genome assembly accession no. GCA_021066465.1). To obtain transcriptome data during spore germination of *A. nidulans*, *A. fumigatus*, and *F. oxysporum*, and mutants of these species, freshly harvested conidia were obtained from CMC medium and cultured on Bird medium at RT (51). Tissue from four morphologically conserved developmental stages [stage 1 (isotropic swelling), Stage 2 (germination), Stage 3 (doubling of the long axis), and Stage 4 (hyphal branching)], were collected from the five species by scraping the surface of colonies from three replicate cultures each. Total RNA was extracted from the collected tissue using TRIzol reagent (Thermo Fisher Scientific, Waltham, MA) according to the manufacturer’s instructions, followed by further purification steps using RNA Clean and Concentrator (Zymo Research Corp., Irvine, CA), and then digestion of genomic DNA with the RNase-Free DNase Set (Qiagen, Hilden, Germany). RNA integrity and quantity were assessed using an Agilent 4200 TapeStation. RNA sequencing libraries were prepared using the Illumina Stranded mRNA Library Kit and sequenced on an Illumina NovaSeq 6000 platform at Michigan State University’s Research Technology Support Facility (https://rtsf.natsci.msu.edu/). The quality of raw reads was assessed using FastQC v0.11.9, and poor-quality reads and adapters were trimmed with Trimmomatic v0.39 (52). Filtered reads were mapped to the reference genomes using HISAT2 v2.1.0 (53), and the resulting BAM files were sorted with SAMtools v1.11 (54). Read counts on genomic features were tallied using HTSeq v0.11.2 (55). The reference genomes were sourced from the Ensembl Fungi database (http://fungi.ensembl.org). RNA-seq was performed on the strains designated as OE::*FoabaA*, OE::*AfabaA*, and *F. oxysporum* WT. RNA was extracted from samples collected at Stage 4 (hyphal branch) in three replicates for each strain. RNA extraction and sequencing were conducted using the same methods as described above.

### Identification of SCOGs and Continuous Ancestral Character Estimation (CACE)

Clusters of orthologous genes were identified using OrthoLoger, a tool associated with OrthoDB, which predicts gene orthologs with default settings (56). Our approach is limited to single-copy genes shared among genomes of the studied species. Tallies of expression such as RPKM are not considered comparable between orthologs among different species, thus, relative gene expression levels across the developmental stages for genes in each species were estimated using LOX (57). We then calculated the fold change of relative expression levels between each adjacent pair of the four developmental stages. The fold change between developmental stages and the phylogeny of the selected species in Newick tree format were supplied as input files to an in-house R program that utilizes the R package APE v5.3, which provided the Continuous Ancestral Character Estimation (58, 59) in expression across adjacent stages at all internal nodes for every SCOG. Expression changes between terminal taxa and the MRCA node were calculated and ranked by the most up-regulated changes toward pathogenesis of interest for further analysis using genetics and bioinformatics tools. For comparison, we examined the FC in expression levels between stages rather than absolute expression levels of reads (17). Values of the FCD were calculated by subtracting the MRCA values from the FC values for the 3,845 SCOGs (**Fig. S1A**). The SCOGs were then ranked according to their FCD values, with the top 10% representing the highly ranked genes and the bottom 10% representing the low ranked genes.

### Differentially expressed genes (DEGs) and principal component analysis (PCA)

The differential expression analysis was performed on the OE::*AfabaA* and OE::*FoabaA* datasets using the DESeq2 R package (v1.30.1). The adjusted p-values (p-adj) > 0.05 and the log fold change (LFC) > 1 were used as thresholds to identify differentially expressed genes (DEGs). We applied principal component analysis (PCA) on the transcriptome dataset to examine the variation in the OE::*AfabaA*, OE::*FoabaA*, and WT strains.

### Targeted *abaA* deletion, swap-out for gene overexpression, and fungal transformation

For the targeted deletion and swap-out of *abaA* and its promoter (**Fig. S9**), *F. oxysporum* was transformed using DNA constructs created through a split marker recombination procedure, as previously described (60). The primers used for amplifying the 5′ and 3′ flanking regions of *abaA* and the final split markers are listed in the Supporting Information **Table S10**. The amplified products were added to fungal protoplasts for transformation, following established protocols (61). For the gene and promoter swap-out, we utilized a geneticin cassette, amplified from the FB009 plasmid (Addgene, Cambridge, MA), as the second selectable marker in *F. oxysporum*. Specifically, for the swap-out of the *ΔFoabaA* strain, transformation was carried out using a DNA construct including *AfabaA* or *FoabaA*, the associated putative promoter region (**Fig. S4, Table S7**), and a selectable marker (geneticin), which was directly added to the protoplasts of *ΔFoabaA*. We obtained the following mutants Δ*FoabaA*, Δ*FoabaA*:: P_AfabaA_:: *AfabaA*, Δ*FoabaA*:: P_FoabaA_:: *AfabaA*, and Δ*FoabaA*:: P_AfabaA_:: *FoabaA*.

### Quantitative PCR, primer, and phylogenetic tree

qPCR analysis was conducted using PowerUp SYBR Green Master Mix (Thermo Fisher Scientific, Waltham, MA). cDNA was synthesized using Maxima™ H Minus cDNA Synthesis Master Mix, with dsDNase (Thermo Fisher Scientific, Waltham, MA) from total RNA. Gene expression levels were evaluated across three biological replicates for each time point. For data normalization, the endogenous control gene EF-1α (FOYG_10039 and AFUA_1G06390) was utilized (62). Primers for PCR and Quantitative PCR (qPCR) were designed using Primer-BLAST from the NCBI site. The primer sequences are listed in Supplementary **Table S10**. Primers were synthesized by Integrated DNA Technologies (IDT, Coralville, IA). A Maximum Likelihood (ML) tree was constructed using the Molecular Evolutionary Genetics Analysis software (MEGA X). This analysis was based on the RNA polymerase II second largest subunit (RPB2) sequences. The ML tree was then inferred with 1,000 bootstraps.

### Scanning electron microscopy

WT and mutant samples cultured on CMC Agar plates for 3 days were used. Each sample was prepared following the method described by Jin et al (63), with the following modifications. Samples were dried using a Leica Microsystems EM CPD300 critical point dryer (Leica Microsystems, Vienna, Austria) with carbon dioxide as the transitional fluid. Dried samples were mounted on aluminum stubs using high vacuum carbon tabs (West Chester, PA). For SEM imaging preparation, the samples were coated with a 15 nm thick osmium layer using a Tennant20 osmium chemical vapor deposition (CVD) coater from Meiwafosis Co., Ltd. (Osaka, Japan). Imaging was performed using a JEOL 7500F scanning electron microscope equipped with a field emission emitter (JEOL Ltd., Tokyo, Japan).

### Pathogenicity tests

To assess pathogenicity, fungal hyphae were inoculated into CMC liquid medium and incubated at 25°C with shaking at 180 rpm for 5 days. Fresh conidia were harvested by filtration through sterilized Miracloth (EMD Millipore Corp., Billerica, MA), followed by centrifugation at 6,000 g for 10 minutes to pellet the conidia. The collected conidia were then diluted to a concentration of 10^5^ conidia/mL in 1x PBS containing 0.1% Tween 80. For infection, *Galleria mellonella* larvae were used as the host model. A 10ul conidial suspension was injected into the hemocoel of each larva through the pro-leg using a sterile 30-gauge syringe. After injection, groups of 20 larvae were placed in sterile 90 mm petri dishes with wood shaving. The larvae were then incubated at 27°C and monitored daily for survival for up to 8 days post-inoculation. Control larvae, also totaling 20 per group, were injected with 10 μL of 1x PBS containing 0.1% Tween 80. Larvae were considered dead when they displayed no movement in response to touch. All experiments were performed with 20 larvae per experimental group with three replicates conducted for each group.

### Pfam domain and HPLS chromosome analysis

Pfam domain analysis and ortholog distribution were conducted using the FungiDB database on the up-regulated DEGs located on the HPLS chromosome in OE::*AfabaA* and OE::*FoabaA*. The distinction between core chromosomes and HPLS chromosomes was based on the findings of Zhang, Yang et al (21). The distribution of Pfam domain orthologs was analyzed at the genus level in fungi, the phylum level in Eukaryota, and the species level in fungi. The analysis included three levels of ortholog identification: genus level in fungi, phylum level in Eukaryota, and species level in fungi. The abbreviations used for specific species are as follows. afum (*A. fumigatus* Af293); anid (*A. nidulans* A4); ncra (*N. crassa* OR74A); focf (*F. oxysporum* f. sp. conglutinans Fo5176); foxr (*F. oxysporum* f. sp. cubense); foxy (*F. oxysporum* f. sp. lycopersici 4287); foxm (*F. oxysporum* f. sp. melonis 26406).

## Accession number(s)

GEO accession no. GSE279549 and BioProject accession no. PRJNA1173228.

## Acknowledgments

We acknowledge the Center for Advanced Microscopy at Michigan State University for providing access to their facilities and technical support.

## Notes

### Competing Interest Statement

The authors have declared no competing interest.

## References

1. Seekles SJ. The breaking of fungal spore dormancy: A coordinated transition. PLOS Biology. 2023;21(4):e3002077.

2. Feofilova EP, Ivashechkin AA, Alekhin AI, Sergeeva YE. Fungal spores: Dormancy, germination, chemical composition, and role in biotechnology (review). Applied Biochemistry and Microbiology. 2012;48(1):1–11.

3. Oh YT, Ahn C-S, Kim JG, Ro H-S, Lee C-W, Kim JW. Proteomic analysis of early phase of conidia germination in Aspergillus nidulans. Fungal Genetics and Biology. 2010;47(3):246–53.

4. Osherov N, May G. Conidial Germination in Aspergillus nidulans Requires RAS Signaling and Protein Synthesis. Genetics. 2000;155(2):647–56.

5. Fillinger S, Chaveroche M-K, van Dijck P, de Vries R, Ruijter G, Thevelein J, et al. Trehalose is required for the acquisition of tolerance to a variety of stresses in the filamentous fungus Aspergillus nidulansThe GenBank accession number for the sequence reported in this paper is AF043230. Microbiology. 2001;147(7):1851–62.

6. Rangel DEN. Stress induced cross-protection against environmental challenges on prokaryotic and eukaryotic microbes. World Journal of Microbiology and Biotechnology. 2011;27(6):1281–96.

7. Rangel DEN, Alder-Rangel A, Dadachova E, Finlay RD, Kupiec M, Dijksterhuis J, et al. Fungal stress biology: a preface to the Fungal Stress Responses special edition. Current Genetics. 2015;61(3):231–8.

8. Wyatt TT, Wösten HAB, Dijksterhuis J. Chapter Two - Fungal Spores for Dispersion in Space and Time. In: Sariaslani S, Gadd GM, editors. Advances in Applied Microbiology. 85: Academic Press; 2013. p. 43-91.

9. Liu F, Zeng M, Zhou X, Huang F, Song Z. Aspergillus fumigatus escape mechanisms from its harsh survival environments. Applied Microbiology and Biotechnology. 2024;108(1):53.

10. Sephton-Clark PCS, Voelz K. Chapter Four - Spore Germination of Pathogenic Filamentous Fungi. In: Sariaslani S, Gadd GM, editors. Advances in Applied Microbiology. 102: Academic Press; 2018. p. 117-57.

11. Baltussen TJH, Zoll J, Verweij PE, Melchers WJG. Molecular Mechanisms of Conidial Germination in Aspergillus spp. Microbiol Mol Biol Rev. 2020;84(1).

12. Ortiz SC, Huang M, Hull CM. Spore Germination as a Target for Antifungal Therapeutics. Antimicrob Agents Chemother. 2019;63(12).

13. Ajmal M, Hussain A, Ali A, Chen H, Lin H. Strategies for Controlling the Sporulation in Fusarium spp. Journal of Fungi [Internet]. 2023; 9(1).

14. Hagiwara D, Takahashi H, Kusuya Y, Kawamoto S, Kamei K, Gonoi T. Comparative transcriptome analysis revealing dormant conidia and germination associated genes in Aspergillus species: an essential role for AtfA in conidial dormancy. BMC Genomics. 2016;17(1):358.

15. Lamarre C, Sokol S, Debeaupuis J-P, Henry C, Lacroix C, Glaser P, et al. Transcriptomic analysis of the exit from dormancy of Aspergillus fumigatus conidia. BMC Genomics. 2008;9(1):417.

16. Wang F, Sethiya P, Hu X, Guo S, Chen Y, Li A, et al. Transcription in fungal conidia before dormancy produces phenotypically variable conidia that maximize survival in different environments. Nature Microbiology. 2021;6(8):1066–81.

17. Trail F, Wang Z, Stefanko K, Cubba C, Townsend JP. The ancestral levels of transcription and the evolution of sexual phenotypes in filamentous fungi. PLOS Genetics. 2017;13(7):e1006867.

18. Kim W, Wang Z, Kim H, Pham K, Tu Y, Townsend Jeffrey P, et al. Transcriptional Divergence Underpinning Sexual Development in the Fungal Class Sordariomycetes. mBio. 2022;13(3):e01100–22.

19. Miguel-Rojas C, Cavinder B, Townsend Jeffrey P, Trail F. Comparative Transcriptomics of Fusarium graminearum and Magnaporthe oryzae Spore Germination Leading up To Infection. mBio. 2023;14(1):e02442–22.

20. Latgé JP. Aspergillus fumigatus and aspergillosis. Clin Microbiol Rev. 1999;12(2):310–50.

21. Zhang Y, Yang H, Turra D, Zhou S, Ayhan DH, DeIulio GA, et al. The genome of opportunistic fungal pathogen Fusarium oxysporum carries a unique set of lineage-specific chromosomes. Communications Biology. 2020;3(1):50.

22. P GR, Kumar A, Tandon R, N HA. Ophthalmic infections caused by Aspergillus nidulans: A case series and short review of literature. Curr Med Mycol. 2021;7(4):43–8.

23. Beadle GW, Tatum EL. Genetic Control of Biochemical Reactions in Neurospora*. Proceedings of the National Academy of Sciences. 1941;27(11):499–506.

24. Davis RH, Perkins DD. Neurospora: a model of model microbes. Nature Reviews Genetics. 2002;3(5):397–403.

25. Gortikov M, Wang Z, Steindorff AS, Grigoriev IV, Druzhinina IS, Townsend JP, et al. Sequencing and Analysis of the Entire Genome of the Mycoparasitic Bioeffector Fungus Trichoderma asperelloides Strain T 203 (Hypocreales). Microbiol Resour Announc. 2022;11(2):e0099521.

26. Ni M, Gao N, Kwon N-J, Shin K-S, Yu J-H. Regulation of Aspergillus Conidiation. Cellular and Molecular Biology of Filamentous Fungi 2010. p. 557–76.

27. Son Y-E, Yu J-H, Park H-S. Regulators of the Asexual Life Cycle of Aspergillus nidulans. Cells. 2023; 12(11).

28. Ruiz-Roldán MC, Köhli M, Roncero MIG, Philippsen P, Di Pietro A, Espeso Eduardo A. Nuclear Dynamics during Germination, Conidiation, and Hyphal Fusion of Fusarium oxysporum. Eukaryotic Cell. 2010;9(8):1216–24.

29. Han K-H, Kim JH, Moon H, Kim S, Lee S-S, Han D-M, et al. The Aspergillus nidulans esdC (early sexual development) gene is necessary for sexual development and is controlled by veA and a heterotrimeric G protein. Fungal Genetics and Biology. 2008;45(3):310–8.

30. Hissen AH, Wan AN, Warwas ML, Pinto LJ, Moore MM. The Aspergillus fumigatus siderophore biosynthetic gene sidA, encoding L-ornithine N5-oxygenase, is required for virulence. Infect Immun. 2005;73(9):5493–503.

31. Haas H. Fungal siderophore metabolism with a focus on Aspergillus fumigatus. Nat Prod Rep. 2014;31(10):1266–76.

32. Schrettl M, Bignell E, Kragl C, Joechl C, Rogers T, Arst HN, Jr., et al. Siderophore biosynthesis but not reductive iron assimilation is essential for Aspergillus fumigatus virulence. J Exp Med. 2004;200(9):1213–9.

33. Ballou ER, Wilson D. The roles of zinc and copper sensing in fungal pathogenesis. Current Opinion in Microbiology. 2016;32:128–34.

34. Eisendle M, Schrettl M, Kragl C, Müller D, Illmer P, Haas H. The Intracellular Siderophore Ferricrocin Is Involved in Iron Storage, Oxidative-Stress Resistance, Germination, and Sexual Development in Aspergillus nidulans. Eukaryotic Cell. 2006;5(10):1596–603.

35. Franken ACW, Lechner BE, Werner ER, Haas H, Lokman BC, Ram AFJ, et al. Genome mining and functional genomics for siderophore production in Aspergillus niger. Briefings in Functional Genomics. 2014;13(6):482–92.

36. Moreno MÁ, Ibrahim-Granet O, Vicentefranqueira R, Amich J, Ave P, Leal F, et al. The regulation of zinc homeostasis by the ZafA transcriptional activator is essential for Aspergillus fumigatus virulence. Molecular Microbiology. 2007;64(5):1182–97.

37. Brown NA, Goldman GH. The contribution of Aspergillus fumigatus stress responses to virulence and antifungal resistance. Journal of Microbiology. 2016;54(3):243–53.

38. Bernard M, Latgé JP. Aspergillus fumigatus cell wall: composition and biosynthesis. Medical Mycology. 2001;39(1):9–17.

39. Frawley D, Karahoda B, Sarikaya Bayram Ö, Bayram Ö. The HamE scaffold positively regulates MpkB phosphorylation to promote development and secondary metabolism in Aspergillus nidulans. Sci Rep. 2018;8(1):16588.

40. Frawley D, Stroe MC, Oakley BR, Heinekamp T, Straßburger M, Fleming AB, et al. The Pheromone Module SteC-MkkB-MpkB-SteD-HamE Regulates Development, Stress Responses and Secondary Metabolism in Aspergillus fumigatus. Frontiers in Microbiology. 2020;11.

41. Johnstone IL, McCabe PC, Greaves P, Gurr SJ, Cole GE, Brow MAD, et al. Isolation and characterisation of the crnA-niiA-niaD gene cluster for nitrate assimilation in Aspergillus nidulans. Gene. 1990;90(2):181–92.

42. Marzluf GA. Regulation of Nitrogen Metabolism in Mycelial Fungi. In: Brambl R, Marzluf GA, editors. Biochemistry and Molecular Biology. Berlin, Heidelberg: Springer Berlin Heidelberg; 1996. p. 357–68.

43. Krappmann S, Braus GH. Nitrogen metabolism of Aspergillus and its role in pathogenicity. Medical Mycology. 2005;43(Supplement_1):S31–S40.

44. Lee IR, Morrow CA, Fraser JA. Nitrogen regulation of virulence in clinically prevalent fungal pathogens. FEMS Microbiology Letters. 2013;345(2):77–84.

45. Filho A, Brancini GTP, de Castro PA, Valero C, Ferreira Filho JA, Silva LP, et al. Aspergillus fumigatus G-Protein Coupled Receptors GprM and GprJ Are Important for the Regulation of the Cell Wall Integrity Pathway, Secondary Metabolite Production, and Virulence. mBio. 2020;11(5).

46. Han K-H, Seo J-A, Yu J-H. Regulators of G-protein signalling in Aspergillus nidulans: RgsA downregulates stress response and stimulates asexual sporulation through attenuation of GanB (Gα) signalling. Molecular Microbiology. 2004;53(2):529–40.

47. Boni AC, Ambrósio DL, Cupertino FB, Montenegro-Montero A, Virgilio S, Freitas FZ, et al. Neurospora crassa developmental control mediated by the FLB-3 transcription factor. Fungal Biology. 2018;122(6):570–82.

48. Mead ME, Borowsky AT, Joehnk B, Steenwyk JL, Shen X-X, Sil A, et al. Recurrent Loss of abaA, a Master Regulator of Asexual Development in Filamentous Fungi, Correlates with Changes in Genomic and Morphological Traits. Genome Biology and Evolution. 2020;12(7):1119–30.

49. O’Donnell K, Gueidan C, Sink S, Johnston PR, Crous PW, Glenn A, et al. A two-locus DNA sequence database for typing plant and human pathogens within the Fusarium oxysporum species complex. Fungal Genetics and Biology. 2009;46(12):936–48.

50. Wang CJ, Thanarut C, Sun PL, Chung WH. Colonization of human opportunistic Fusarium oxysporum (HOFo) isolates in tomato and cucumber tissues assessed by a specific molecular marker. PLoS One. 2020;15(6):e0234517.

51. Metzenberg RL. Bird Medium: an alternative to Vogel Medium. Fungal Genetics Reports. 2004(51(1)):19-20.

52. Bolger AM, Lohse M, Usadel B. Trimmomatic: a flexible trimmer for Illumina sequence data. Bioinformatics. 2014;30(15):2114–20.

53. Kim D, Langmead B, Salzberg SL. HISAT: a fast spliced aligner with low memory requirements. Nat Methods. 2015;12(4):357–60.

54. Li H, Handsaker B, Wysoker A, Fennell T, Ruan J, Homer N, et al. The Sequence Alignment/Map format and SAMtools. Bioinformatics. 2009;25(16):2078–9.

55. Anders S, Pyl PT, Huber W. HTSeq--a Python framework to work with high-throughput sequencing data. Bioinformatics. 2015;31(2):166–9.

56. Kuznetsov D, Tegenfeldt F, Manni M, Seppey M, Berkeley M, Kriventseva Evgenia V, et al. OrthoDB v11: annotation of orthologs in the widest sampling of organismal diversity. Nucleic Acids Research. 2023;51(D1):D445–D51.

57. Zhang Z, López-Giráldez F, Townsend JP. LOX: inferring Level Of eXpression from diverse methods of census sequencing. Bioinformatics. 2010;26(15):1918–9.

58. Schluter D, Price T, Mooers AØ, Ludwig D. Likelihood of ancestor states in adaptive radiation. Evolution. 1997;51(6):1699–711.

59. Paradis E, Claude J, Strimmer K. APE: analyses of phylogenetics and evolution in R language. Bioinformatics. 2004;20(2):289–90.

60. Kim H-K, Lee S, Jo S-M, McCormick SP, Butchko RAE, Proctor RH, et al. Functional Roles of FgLaeA in Controlling Secondary Metabolism, Sexual Development, and Virulence in Fusarium graminearum. PLOS ONE. 2013;8(7):e68441.

61. Kim H-K, Jo S-M, Kim G-Y, Kim D-W, Kim Y-K, Yun S-H. A Large-Scale Functional Analysis of Putative Target Genes of Mating-Type Loci Provides Insight into the Regulation of Sexual Development of the Cereal Pathogen Fusarium graminearum. PLOS Genetics. 2015;11(9):e1005486.

62. Kozera B, Rapacz M. Reference genes in real-time PCR. Journal of Applied Genetics. 2013;54(4):391–406.

63. Fischer ER, Hansen BT, Nair V, Hoyt FH, Dorward DW. Scanning electron microscopy. Curr Protoc Microbiol. 2012;Chapter 2:Unit 2B.

